# Determination of GLP-1 Secretion Potential of Dead and Live *Akkermansia muciniphila* Using Human L-cells

**DOI:** 10.64898/2026.03.18.708496

**Authors:** Subhendu Nayak, Prabhu Rajagopalan, Raksha Sunhare, Shalini Jain

## Abstract

**Background/Objectives:** Glucagon-Like Peptide-1 (GLP-1) is a key incretin hormone that regulates glucose homeostasis and energy metabolism. Impaired GLP-1 signaling contributes to the development of obesity, metabolic syndrome, and type 2 diabetes. Emerging evidence indicates that gut microbiota-derived components can influence GLP-1 secretion, highlighting the therapeutic potential of microbial modulators. *Akkermansia muciniphila*, a next-generation probiotic associated with improved metabolic health, remains underexplored for its capacity to stimulate GLP-1 release. This study aimed to investigate the GLP-1– stimulatory effects of live and pasteurized (dead) *A. muciniphila* strains in human enteroendocrine cells.

**Methods:** Human enteroendocrine L-cells (NCI-H716) were treated with varying doses of live and dead *A. muciniphila* from Vidya Herbs’s proprietary VHAKM strain and a commercially available marketed strain (dead form). Following incubation, GLP-1 levels were quantified from culture supernatants using enzyme-linked immunosorbent assay (ELISA). Comparative analyses assessed differences in GLP-1 secretion between strains and treatment forms.

**Results:** Both live and pasteurized VHAKM strains significantly increased GLP-1 secretion compared to untreated controls. The live VHAKM strain exhibited higher GLP-1 stimulatory activity than its pasteurized counterpart and the marketed strain. The results suggest a strain-specific and viability-dependent modulation of GLP-1 secretion in human L-cells.

**Conclusions:** This study demonstrates that *A. muciniphila* VHAKM enhances GLP-1 secretion in a strain- and form-dependent manner, with live cells showing superior efficacy. These findings provide foundational insights for developing microbiome-targeted interventions to boost endogenous GLP-1 levels and improve metabolic health outcomes.

## 1. Introduction

Glucagon-Like Peptide-1 (GLP-1) is a key incretin hormone secreted by enteroendocrine L-cells in the gut, playing a pivotal role in regulating postprandial insulin secretion, glucose homeostasis, appetite suppression, and gastric emptying [1]. GLP-1-based therapies, such as GLP-1 receptor agonists (GLP-1RAs), including semaglutide (Ozempic, Wegovy), liraglutide (Victoza, Saxenda), and dulaglutide (Trulicity), have revolutionized the management of type 2 diabetes and obesity. However, the rising prevalence of metabolic disorders—affecting over 650 million adults globally and incurring healthcare costs exceeding $200 billion annually in the United States alone[2]—underscores the urgent need for novel and alternative strategies. Despite their clinical efficacy, GLP-1RAs are associated with high costs, gastrointestinal side effects, and limited accessibility, which constrain their broader utilization. Therefore, alternative approaches to stimulate endogenous GLP-1 secretion using safe and cost-effective modalities are critically needed. The gut microbiome has emerged as a central player in host metabolism [3-5], with increasing evidence highlighting its role in the pathogenesis of obesity, insulin resistance, and metabolic syndrome. Microbiome-derived metabolites, such as short-chain fatty acids (SCFAs), indole derivatives, and secondary bile acids, have been shown to influence GLP-1 secretion through enteroendocrine signaling pathways [6]. For instance, SCFAs like acetate and butyrate stimulate GLP-1 via G-protein coupled receptors (GPR41 and GPR43) [7]. Furthermore, bacterial components such as lipopolysaccharides (LPS) and peptidoglycans can modulate enteroendocrine functions through Toll-like receptor pathways [8]. These findings position the gut microbiome as a promising target for modulating GLP-1 physiology, providing a basis for exploring microbial interventions to enhance endogenous GLP-1 levels.

*Akkermansia (A*.*) muciniphila*, a mucin-degrading bacterium residing in the mucus layer of the gut, has gained prominence as a next-generation probiotic due to its beneficial effects on metabolic health.[9] Multiple preclinical and clinical studies have demonstrated *Akkermansia*’s role in improving glucose tolerance, reducing adiposity, and enhancing gut barrier integrity [10]. Live *A. muciniphila* exerts its beneficial effects primarily through its ability to degrade host-derived mucin in the intestinal mucus layer, a process that not only supports its own growth but also shapes the gut microenvironment. During mucin degradation, *A. muciniphila* secretes specific outer membrane proteins, notably P9, which interact with host epithelial immune cells, and enteroendocrine L-cells [11]. L cells are a special type of cells that when given information to do so can secrete GLP-1 [12]. L-cells are in the ileum and colon, which are highly absorbent areas for the bloodstream in the intestinal tract. Therefore, when P9 stimulates L-cells, GLP-1 can enter the bloodstream and head to the pancreas [11]. Once GLP-1 is present in the pancreas, it can then trigger it to release insulin [13].

Live *A. muciniphila* functions as a keystone species for maintaining gut homeostasis and systemic metabolic balance through mucin degradation.[9] Notably, live forms of *A. muciniphila* have shown metabolic benefits, while pasteurized forms only contain metabolites and their effects. However, strain-dependent variations in bioactivity are increasingly recognized, and direct comparisons of live versus dead forms regarding GLP-1 stimulation remain scarce. Systematic evaluations of commercial and proprietary *A. muciniphila* strains are limited, necessitating well-designed studies to unravel their functional capacities and translational relevance for GLP-1-centric metabolic interventions.

## 2. Materials and Methods

### Cell Culture and Conditions

Human enteroendocrine NCI-H716 cells were cultured in RPMI-1640 medium supplemented with 10% fetal bovine serum (FBS), 1% L-glutamine, and antibiotics (100 U/mL penicillin and 100 µg/mL streptomycin). Cells were maintained at 37°C in a humidified atmosphere containing 5% CO2 and were passaged every 3–4 days to maintain exponential growth.

### Culture of A. muciniphila Strains

*Akkermansia muciniphila VHAKM was isolated from human fecal material. Its potential probiotic properties are characterized by functional genome analysis. Akkermansia muciniphila VHAKM’s entire genome sequence will enable researchers to fully understand the molecular specifics of the organism’s features and safety. The percentage of identity is more than 99% between Vidya and ATCC BAA-835 strain; and the whole genome sequence can be found on:*

*https://www.ncbi.nlm.nih.gov/nucleotide/PV774913.1?report=genbank&log$=nuclalign&blast_rank=1&RID=5R03TD43013*.

*A. muciniphila* VHAKM strain (Vidya Herbs) and Marketed (Mrkt) AKMS strain (https://pendulumlife.com/) were cultured anaerobically in brain heart infusion (BHI) broth supplemented with 0.5% porcine mucin. Cultures were incubated at 37°C for 48 hours under anaerobic conditions using an anaerobic chamber with a gas mixture of 5% CO2, 5% H2, and 90% N2. For pasteurization, bacterial cultures were subjected to heat treatment at 70°C for 30 minutes to obtain dead cells. Furthermore, dead cells were disrupted by homogenization protocol and homogenate was further centrifuged and clear supernatants were used for cell treatments. Total protein was measured using BCA method.

### GLP-1 Secretion Assay in NCI-H716 Cells

The NCI-H716 cells were seeded in 24-well plates at a density of 2 × 10^5 cells/well and allowed them to adhere overnight and let them become around 70% confluent. Cells were washed with PBS and incubated in serum-free medium for 2 hours prior to treatment. Live and pasteurized *A. muciniphila* cells were added at concentrations of 1.25, 2.5, 5 and 10 billion CFUs/ mL as indicated in table here. After 30 and 60 minutes of incubation, culture supernatants were collected and centrifuged to remove debris. GLP-1 levels were quantified using a human GLP-1 (active) ELISA kit (MilliporeSigma), following the manufacturer’s instructions.

**Table 1.**
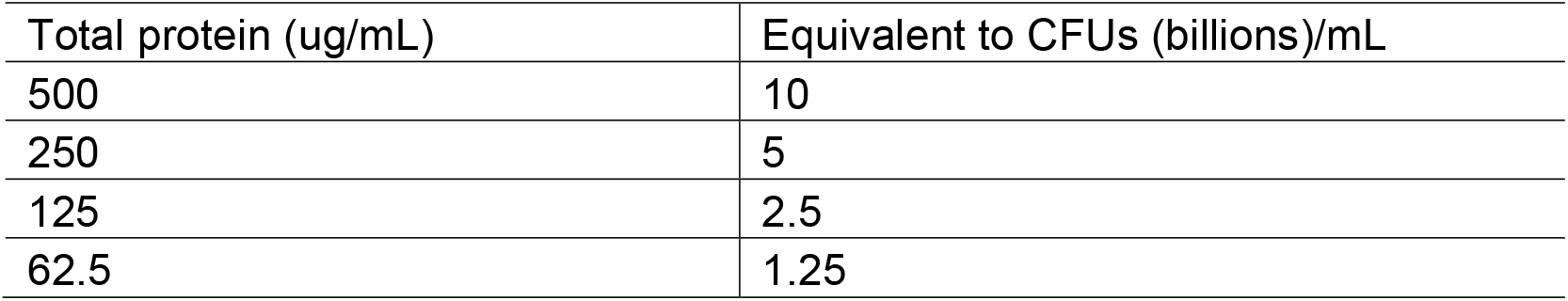
Doses of bacterial supernatants.

| **Table 1. Doses of bacterial supernatants** |  |
| --- | --- |
| Total protein (ug/mL) | Equivalent to CFUs (billions)/mL |
| 500 | 10 |
| 250 | 5 |
| 125 | 2.5 |
| 62.5 | 1.25 |

### Comparative Studies of Live and Dead Forms

To evaluate the differential effects of live versus pasteurized forms, parallel treatments with live and dead VHAKM as well as dead VHAKM versus dead MrktAKMS strains were performed under identical conditions. Cell viability assays (MTT assay) were conducted to confirm no cytotoxicity at tested concentrations. Negative control (no treatment) and positive control (10 mM glutamine) were included in each experiment.

### Data Analysis

All experiments were performed in triplicates and repeated independently at least three times. Data are presented as mean ± SEM. Statistical analyses were conducted using MS Excel and GraphPad Prism 9. Differences between groups were assessed using unpaired two-tailed Student’s t-test or one-way ANOVA followed by Tukey’s post hoc test where applicable. A p-value of <0.05 was considered statistically significant.

## 3. Results

### Both live and dead VHAKM cells stimulate GLP-1 secretion from human L-cells

Our earlier studies show that the pasteurized (dead) VHAKM cells increase GLP-1 secretion in NCI-H716 L-cells in a dose-dependent manner,[14] however, it was unknown whether live and dead VHAKM show similar or distinct effects on GLP-1 secretion. Our current studies demonstrated that both live and dead significantly increased GLP-1 secretion from human L-cells in dose dependent manner (Figure 1). At 10^9 CFU/mL, GLP-1 levels were elevated by 1.8-fold compared to the untreated control (p < 0.01; Figure 1A). Lower doses (10^7 and 10^8 CFU/mL) also showed significant increases (p < 0.05), indicating that live and dead VHAKM cells exhibit bioactivity for GLP-1 stimulation (Figure 1).

**Figure 1.**
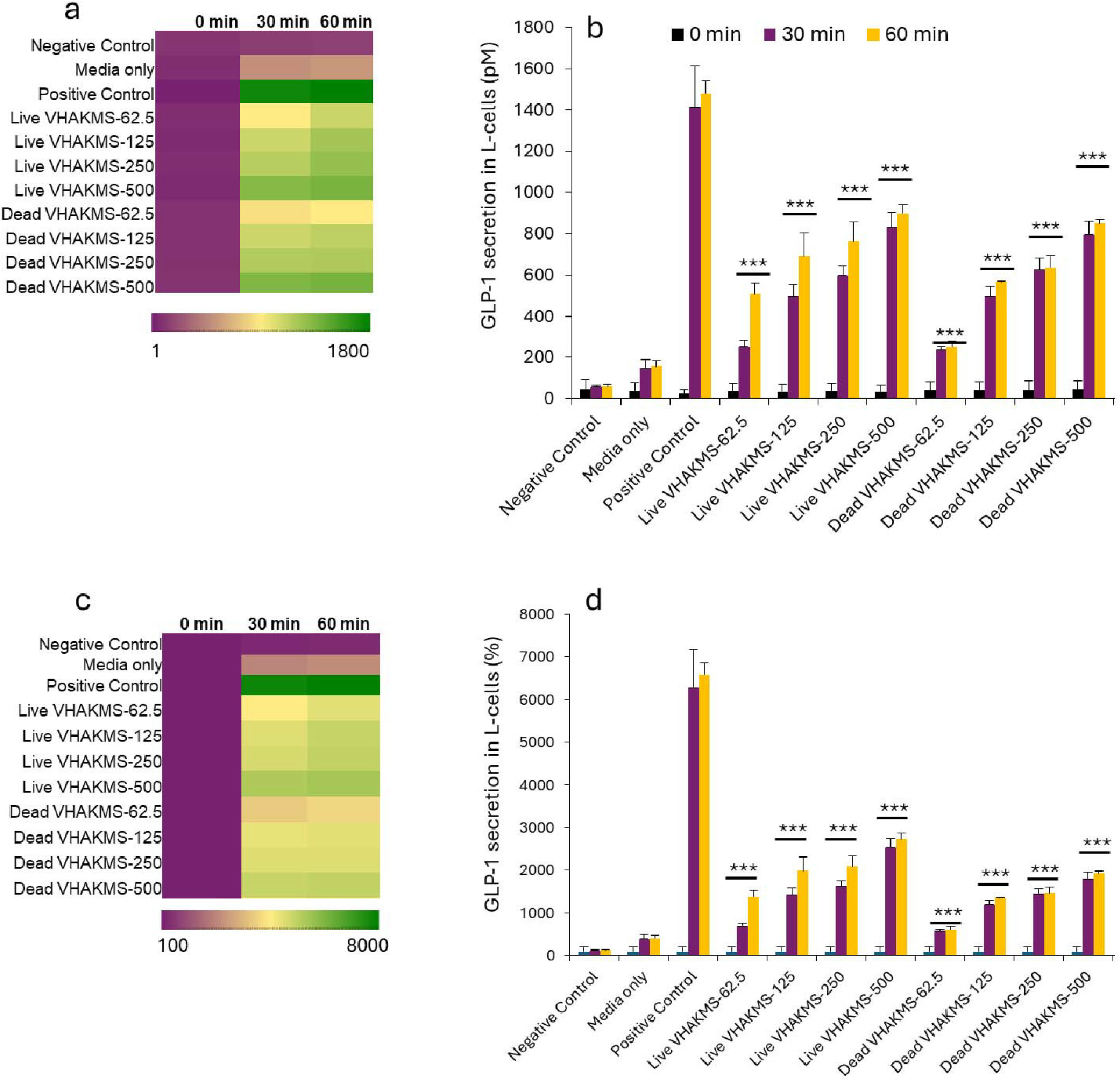
Both live and heat-killed AKMS stimulate GLP-1 secretion from human L-cells. (a, b) Heatmap (a) and bar graphs (b) show that both live and heat-killed (dead) AKMS stimulate GLP-1 secretion (presented as absolute GLP-1 levels in pM) in a dose- and time-dependent manner from human NCI-H716 L-cells. Similarly, the percent increase in GLP-1 secretion is shown in the heatmap (c) and bar graphs (d) for both live and heat-killed (dead) AKMS in a dose- and time-dependent manner. All values presented in these graphs represent the mean ± standard error of the mean from experiments performed in triplicate. Values marked with ***P < 0.001 indicate statistical significance.

### Live VHAKM is slightly more effective in stimulating GLP-1 than its heat-killed form

To further evaluate potential differences in the GLP-1 stimulatory activity between live and heat-killed VHAKM, we performed a comparative analysis using human NCI-H716 L-cells. The results demonstrated that live VHAKM induced a greater secretion of GLP-1 compared to its heat-killed (pasteurized) counterpart (Figure 2). The percent change in GLP-1 secretion relative to baseline controls showed a markedly higher response with the live strain specifically at higher dose of 500, suggesting that bacterial viability contributes to the observed bioactivity.

**Figure 2.**
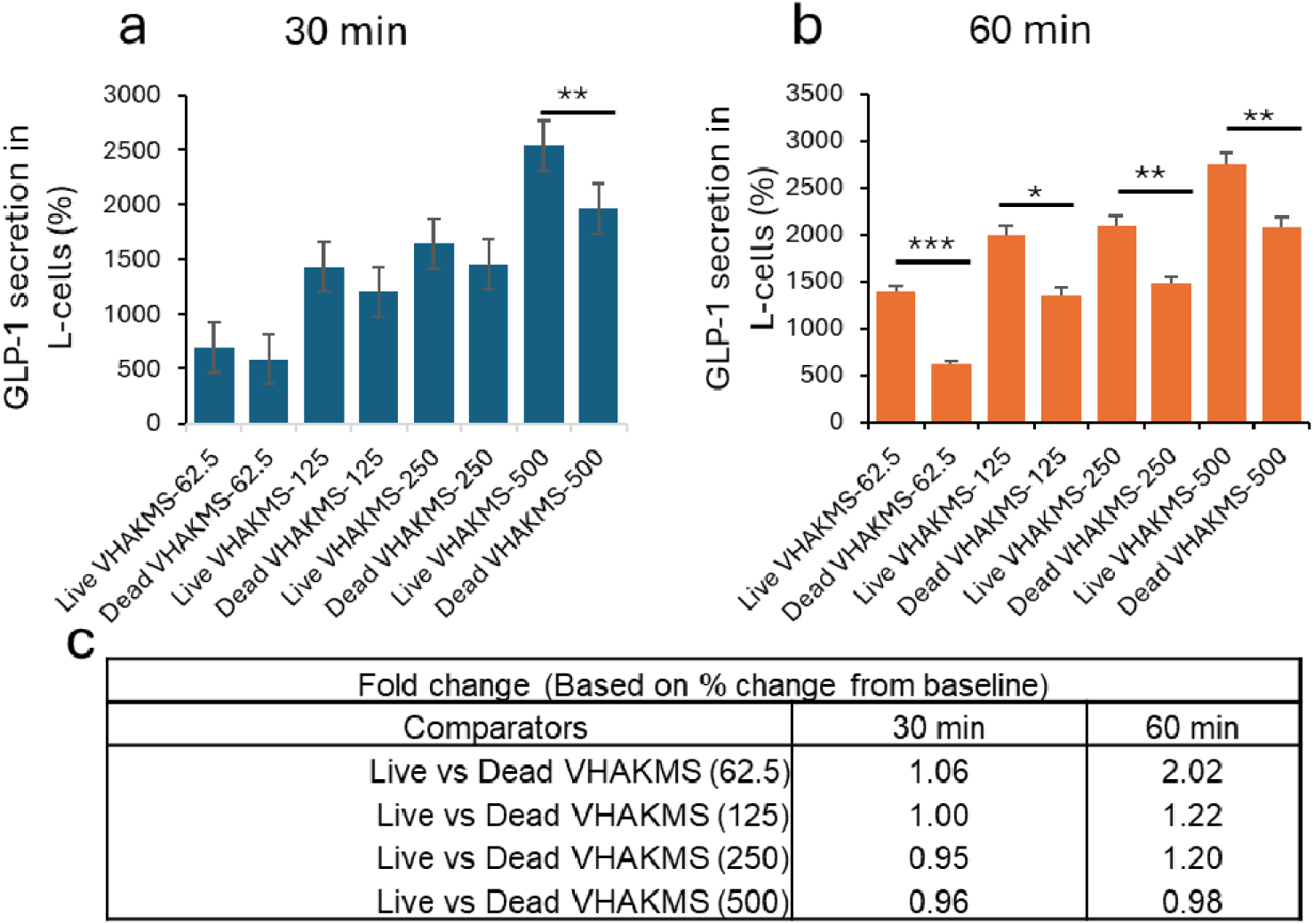
Live VHAK show significantly greater GLP-1 stimulatory effects than their heat-killed counterparts. Upper panels (a and b) show the absolute GLP-1 secretion levels from human L-cells after 30 (a) and 60 (b) minutes of treatment, while lower panels (c and d) depict the percent (%) change in GLP-1 secretion from human L-cells after 30 (c) and 60 (d) minutes of treatment with live and heat-killed VHAK. (e) The table presents the fold change in GLP-1 secretion by live versus heat-killed VHAKSin human L-cells. All values are presented as means, with error bars representing the standard error of the mean (SEM). Statistical significance is indicated as *p < 0.05; **p < 0.01; and ***p < 0.001.

**Figure 3.**
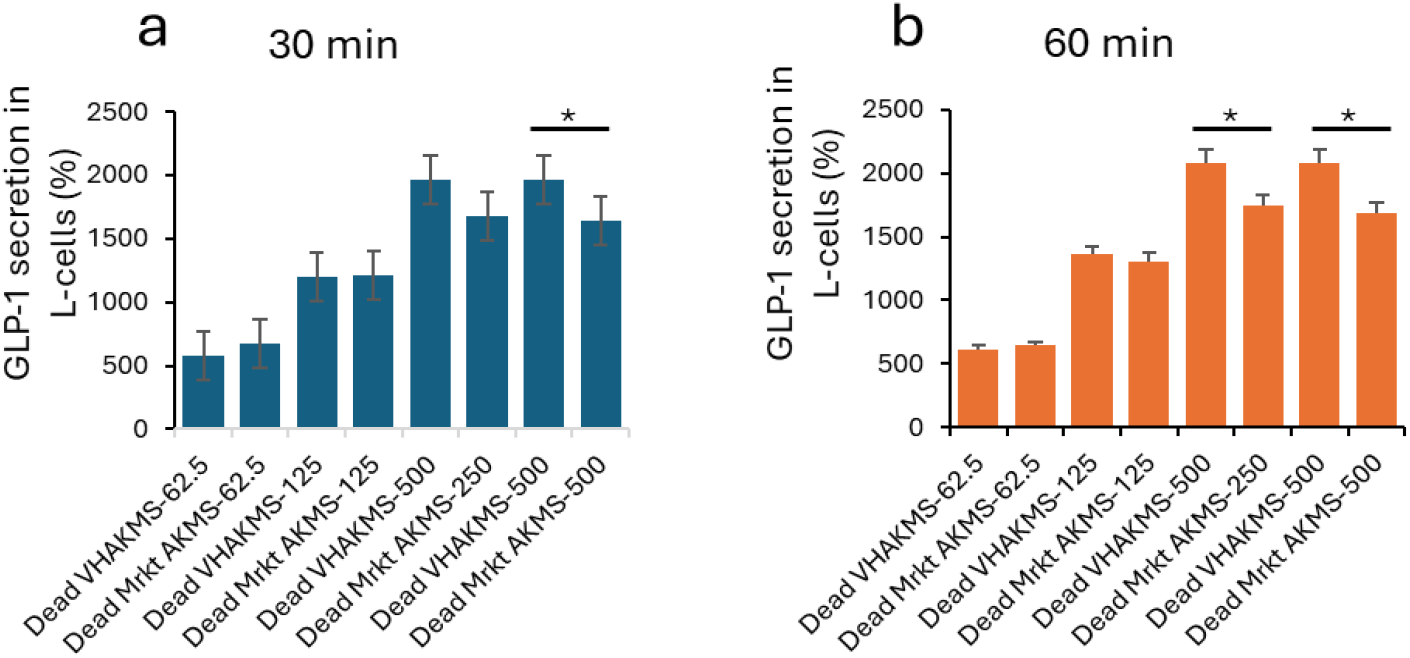
Heat-killed VHAKM shows slightly greater GLP-1 stimulatory activity than heat-killed marketed AKMS (MrktAKMS). Differences in GLP-1 secretion between heat-killed Vidya Herbs AKMS (VHAKM) and heat-killed marketed AKMS (MrktAKMS) are shown as percent (%) changes in GLP-1 secretion from human L-cells after 30 (c) and 60 (d) minutes of treatment. Bar graph values represent the mean, with error bars indicating the standard error of the mean (SEM). Statistical significance is denoted as *p < 0.05; **p < 0.01; and ***p < 0.001.

Quantitatively, live VHAKM exhibited approximately 1.2-to 2-fold higher GLP-1 stimulatory activity than the heat-killed strain, with the most pronounced effect observed after 60 minutes of incubation (p < 0.001; Figure 2e). These findings indicate that while both live and heat-killed VHAKM can stimulate GLP-1 release, the live form retains superior bioactivity, potentially due to the presence of metabolically active components or secreted factors that enhance enteroendocrine cell signaling.

### Both Dead VHAKM and MrktAKMS Strains Effectively Stimulate GLP-1 Secretion

Both dead VHAKM and the MrktAKMS strains significantly enhanced GLP-1 secretion in human NCI-H716 L-cells. At a concentration of 10 billion CFU/mL, heat-killed VHAKM cells induced a slightly higher increase in GLP-1 secretion (19.7% [1966.07 vs 1641.12]) compared to heat-killed MrktAKMS (p < 0.05; Figure 5a) after 30 minutes of incubation. Interestingly, after 60 minutes of incubation, both 5 billion and 10 billion CFU/mL doses of heat-killed VHAKM and MrktAKMS more robustly increased GLP-1 secretion (19.1 % [2078.6 vs 1744] and 23.3% [2077.56 vs 16.85.3], respectively) compared to baseline controls (Figure 5b). These findings indicate that even in their non-viable (pasteurized) form, both strains retain GLP-1 stimulatory activity, with VHAKM showing slightly greater efficacy than the marketed strain. No cytotoxic effects were observed at any of the tested doses, confirming the safety of these bacterial preparations under experimental conditions.

## 4. Discussion

Our study provides several salient discoveries. First, we show that both live and heat-killed (pasteurized) *A. muciniphila* (VHAKM strain) cells stimulate GLP-1 secretion in human NCI-H716 L-cells in a dose- and time-dependent manner. Second, when comparing live versus heat-killed (dead) VHAKM cells, the live cells induced modestly but significantly greater GLP-1 secretion. Third, we compared VHAKM strain to a marketed AKMS strain (MrktAKMS), both in the heat-killed form, and found that while both stimulated GLP-1 secretion, VHAKM achieved slightly higher efficacy. Finally, no cytotoxicity was observed at the doses tested, supporting safety of the preparations under in vitro conditions.

In interpreting these findings, it is useful to place them in the context of prior literature. The capacity of *A. muciniphila* to improve metabolic health (insulin sensitivity, adiposity reduction, gut barrier integrity) is now well-established [15, 16]. More specifically regarding GLP-1 secretion, a landmark study by Plovier and colleagues identified a secreted protein “P9” from A. muciniphila that binds ICAM-2 on enteroendocrine cells, triggers PLC/Ca^2+^/CREB signaling, and increases GLP-1 secretion in vitro and in high-fat-diet mice [11, 17]. That study points to viable metabolic-signaling activity of live (or at least metabolically active) bacteria or their secreted components. Other recent reviews indicate that *A. muciniphila* (and its outer membrane protein Amuc_1100) may promote GLP-1 secretion, improve gut barrier and metabolic outcomes [18].

Our results align with and extend those observations in several important ways. First, we provide direct in vitro evidence that both live and non-viable (heat-killed) cells of a defined *A. muciniphila* strain stimulate GLP-1 secretion in a human L-cell model. This suggests that viability per se is not strictly required for GLP-1 stimulatory bioactivity — consistent with several reports of “postbiotic” activity of pasteurized *A. muciniphila* showing metabolic benefits in animal models [17]. Second, our comparative live vs dead data add nuance: although the heat-killed cells are active, live cells show modest superior potency. This suggests that some viability-dependent mechanisms (e.g., metabolite secretion, cell surface molecule maintenance, dynamic interactions) may augment the effect, which is an insight not previously quantified in many studies. Third, the strain-comparison (VHAKM vs MrktAKMS) underscores the importance of strain specificity in probiotic/postbiotic effects — consistent with the overall literature emphasizing that not all strains of *A. muciniphila* are equivalent in effect. Our finding that VHAKM slightly outperformed a marketed strain suggests possible proprietary enhancements (or simply strain variation) in GLP-1 stimulating capacity.

At the same time, our data diverge in some respects from prior reports and highlight knowledge gaps. For example, in some animal studies pasteurized *A. muciniphila* was reported to outperform live bacteria in metabolic endpoints (e.g., weight gain, insulin resistance) whereas our results show the live form had modestly better GLP-1 induction. This raises interesting mechanistic questions: perhaps for whole-body metabolic outcomes the non-viable preparation exerts optimal effect, but for pure GLP-1 secretion the live form retains more potency. Additionally, while the P9 protein story suggests a defined secreted molecule is sufficient to drive GLP-1 release, our data suggest that “whole-cell” preparations (live and dead) can have relatively rapid (30 and 60 min) effects on human enteroendocrine cells in vitro. This raises the possibility of multiple mechanisms: cell surface molecules, released vesicles or metabolites, membrane proteins, or residual enzymatic activity in heat-killed cells. The area of “postbiotic” mechanisms remains under-explored [19].

Mechanistically, how might *A. muciniphila* (live or dead) stimulate GLP-1 secretion? Potential pathways include: (i) the release or presentation of bacterial proteins (e.g., Amuc_1100, P9) that engage enteroendocrine cell receptors such as ICAM-2, triggering intracellular Ca^2+^, PLC and CREB signaling for GLP-1 gene expression/secretion [20]. (ii) Bacterial derived short-chain fatty acids (SCFAs) and bile acid metabolites produced by viable bacteria that engage G-protein-coupled receptors (GPR41/43, TGR5) on L-cells to stimulate hormone release — this is part of the broader gut–metabolite–enteroendocrine axis described in reviews [16]. (iii) Structural cell components (lipids, outer membrane proteins, vesicles) of bacteria that persist after pasteurization and can engage host pathways. The observation that heat-killed cells retain activity supports mechanism (iii). Our finding that live cells are modestly superior supports a role for viability-dependent factors (ii) and (i). Future mechanistic experiments are required to dissect relative contributions of these pathways.

From a translational perspective, our results have several implications. The fact that heat-killed cells retain GLP-1 stimulatory activity simplifies regulatory and safety considerations for development of postbiotic interventions — heat inactivation avoids live-bacterium regulatory concerns and may improve shelf stability. The superiority (albeit modest) of live cells indicates there may still be benefit to live formulations if they can be safely delivered. The strain-specific data supports the need for rigorous strain characterization and selection for optimized bioactivity rather than treating all *A. muciniphila* strains interchangeably. Given the central role of GLP-1 in metabolic regulation (insulin secretion, appetite, gastric motility), our findings support the further exploration of *A. muciniphila* preparations as adjuncts in metabolic disease (obesity, type 2 diabetes) interventions.

However, limitations of our study must be acknowledged. The experiments were conducted in vitro using a human L-cell line (NCI-H716) and may not fully replicate the complexity of the in vivo gut environment (microbiome interactions, gut barrier, immune signaling, endocrine axis, digestion/metabolism). We assessed only short incubation time points (30, 60 min) and did not examine longer time points or repeated exposure. We also did not dissect the mechanistic basis of GLP-1 stimulation (e.g., receptor engagement, signaling pathways, metabolite production) nor examine downstream functional effects (insulin secretion, glucose tolerance). The doses tested (CFU equivalent) may not directly translate to in vivo exposure or deliverable human dose. Additionally, although we compared one marketed strain, broader strain-comparative work would strengthen generalizability. The modest difference between live and heat-killed raises questions of cost-benefit for live formulations. Finally, cell culture models do not account for host absorption, DPP-4 degradation of GLP-1, gut transit time or immunologic context (DPP-4 cleavage reduces circulating GLP-1– only ∼10–15% of secreted GLP-1 reaches the circulation intact).

## 5. Conclusions

In summary, this study demonstrates that both live and heat-killed VHAKM (A. muciniphila) cells significantly stimulate GLP-1 secretion in human L-cells in a dose- and time-dependent manner. The live formulation offers modestly greater efficacy than the heat-killed form, and the proprietary VHAKM strain exhibits slightly superior activity compared to a marketed AKMS strain. These findings highlight the potential of both probiotic and postbiotic A. muciniphila preparations to activate enteroendocrine GLP-1 pathways — with implications for metabolic disease management via incretin modulation. The translational promise is bolstered by the retained bioactivity of non-viable cells (postbiotics), which may simplify regulatory and formulation challenges while preserving effect. Future in vivo and mechanistic studies are warranted to validate efficacy, explore mechanism, optimize dosing and strain, and ultimately move toward human clinical application.

## Supplementary Materials

No supplementary information is provided related to this manuscript.

## Author Contributions

Conceptualization, SN, PR, RS and SJ; methodology, SN, SJ.; formal analysis, SJ.; writing—review and editing, SN, SJ visualization, SJ.; supervision, SJ.; project administration, SN, SJ,; funding acquisition, SN and SJ. All authors have read and agreed to the published version of the manuscript.

## Institutional Review Board Statement

Not applicable.

## Informed Consent Statement

Not applicable.

## Data Availability Statement

Data will be made available upon reasonable request by authors.

## Acknowledgments

Authors are thankful for Vidya Herbs USA, MusB Research Team for their unwavering support and great quality work.

## Abbreviations

The following abbreviations are used in this manuscript:

GLP-1: Glucagon-Like Peptide-1
GLP-1RAs: Glucagon-Like Peptide-1 Receptor Agonists
ELISA: Enzyme-linked immunosorbent assay
VHAKM: Vidya Herbs *Akkermansia muciniphila*
MrktAKMS: Marketed *Akkermansia muciniphila*
GPR: G-protein coupled receptors
LPS: Lipopolysaccharides
CFU: Colony Forming Unit
H2: Hydrogen
N2: Nitrogen

